# Sleep deprivation, cytokine dysregulation, and the risk of cardiac arrhythmia in domestic dogs

**DOI:** 10.64898/2025.12.23.696165

**Authors:** Adithya Sethumadhavan, Arun HS Kumar

## Abstract

**Background:** Sleep deprivation is increasingly recognized as a potent physiological stressor capable of altering immune function and promoting systemic inflammation. Emerging evidence suggests that these inflammatory changes may contribute to cardiac electrical instability and arrhythmia risk. However, the specific cytokines involved and their mechanistic pathways in domestic dogs remain poorly characterized. This study aimed to (1) identify key cytokines consistently associated with sleep deprivation through a systematic meta-analysis of the literature, and (2) perform a domestic dog-specific network and Gene Ontology enrichment analysis to characterize their interactions and potential mechanistic links to arrhythmogenesis.

**Methods:** A structured search of PubMed and Google Scholar identified 25 articles, of which 24 unique records were screened and seven met all inclusion criteria. From these studies, seven cytokines (IL-6, IL-17A, TNF, IL-1, IL-21, IFN-γ, and CRP) were consistently associated with sleep deprivation and were subjected to domestic dog-specific protein-protein interaction analysis using the STRING database. Network topology and enriched biological processes were evaluated to identify mechanistic pathways connected to cardiac electrophysiology.

**Results:** Network analysis revealed several highly interconnected signalling nodes, including IL-10, IL-1β, JAK1, JAK2, IL-6, STAT3, TNF, IFN-γ, IL-2, and IL-4. Enrichment analysis indicated activation of processes such as chemokine production, positive regulation of osteoclast differentiation, JAK-STAT signalling, membrane protein ectodomain proteolysis, and regulation of vitamin D metabolism. Integration of primary and secondary networks revealed three major clusters cantered on interleukin signalling, TNF-driven pathways, and CRP-associated acute-phase responses. Collectively, these pathways converge on cytokine-mediated mechanisms known to influence myocardial conduction and increase arrhythmia susceptibility.

**Conclusions:** These findings suggest that sleep deprivation activates coordinated inflammatory networks that may predispose dogs to arrhythmias through multiple interconnected biological pathways. Therapeutic strategies addressing both inflammation and electrical instability may be required to fully manage arrhythmia risk in sleep-deprived canine patients.

## Introduction

In veterinary medicine, there is growing recognition that sleep quality influences cardiovascular health in dogs.^[1-3]^ Much of the current evidence derives from studies of brachycephalic breeds with Brachycephalic Obstructive Airway Syndrome (BOAS), where sleep-disordered breathing leads to chronic sleep disruption.^[4,5]^ These dogs demonstrate features like human obstructive sleep apnoea, including hypoxemia, autonomic nervous system (ANS) imbalance, and increased arrhythmia risk.^[4,5]^ However, beyond autonomic dysregulation, emerging research indicates that sleep deprivation (SD) induces profound changes in immune signalling, which may contribute to arrhythmogenesis in canine patients.^[6-9]^ Several experimental studies in humans and laboratory models have shown that SD is associated with the upregulation of pro-inflammatory cytokines, particularly interleukin-6 (IL-6) and interleukin-17A (IL-17A).^[10-12]^ These mediators are central to the initiation of cytokine-driven inflammatory cascades. IL-6, for example, has been reported to increase more than threefold at the transcriptional level following acute sleep loss, with TNF-α messenger RNA levels doubling under the same conditions.^[10,11]^ Partial sleep deprivation also enhances the ability of monocytes to express IL-6 and TNF-α immediately after disrupted sleep,^[13-15]^ highlighting how even modest reductions in sleep duration can trigger exaggerated immune activation. If extrapolated to canine physiology, these findings are highly relevant. The heart, as a continuously active muscle, relies on restorative processes during sleep to repair daily stress-induced damage.^[16,17]^ Cytokines normally play a controlled role in muscle repair and immune regulation; however, chronic elevations of pro-inflammatory cytokines due to insufficient sleep may prevent resolution of the inflammatory response.^[18,19]^ Over time, this persistent low-grade inflammation could compromise myocardial tissue integrity, disrupt ion channel function, and alter gap junction communication, thereby creating suitable conditions for arrhythmia development.^[20,21]^

SD is known to prime innate immune cells such as neutrophils and monocytes, leading to transcriptional reprogramming and increased release of inflammatory mediators.^[22,23]^ In dogs, this would manifest as systemic inflammation, endothelial dysfunction, and impaired vasodilatory responses, all of which burden the cardiovascular system.^[7,23]^ When combined with the sympatho-vagal coactivation observed in BOAS^[5]^ and other sleep-disrupted states, the consequence is a synergistic increase in arrhythmia susceptibility. Adrenergic surges, particularly upon arousal from fragmented sleep, further amplify inflammatory mediator release, accelerating this cycle of inflammation and electrophysiological instability.^[24,25]^ Hence, the interplay between sleep deprivation, cytokine dysregulation, and autonomic imbalance presents a compelling mechanistic framework for arrhythmia risk in dogs. Pro-inflammatory cytokines such as IL-6 and TNF-α not only perpetuate systemic inflammation but also exert direct effects on cardiac electrical activity, while ANS dysregulation lowers the threshold for arrhythmia initiation.^[26,27]^ Cytokines play a vital role in regulating the immune system by activating mechanisms to counter and mitigate cell death, antigens, and inflammation.^[28,29]^ Once the pharmacology of cytokines is achieved, they are cleared from the body through natural elimination or anti-inflammatory cytokines. However, an overproduction of cytokines can disrupt a balanced immune system and begin to cause damage to surrounding cells. Sleep duration and quality is essential for the body’s ability to repair itself. If sleep quality/duration is impaired, it can leave the body in a partial stage of repair, resulting in the build-up of pro-inflammatory cytokines produced by the immune system. A buildup of such cytokines can render the system, and in this case the heart, in a constant state of inflammation and reduced immunity. For breeds predisposed to sleep-disordered breathing, these processes may underlie the increased frequency of life-threatening arrhythmias observed clinically.^[4,5,30]^ Further investigation into cytokine dynamics in canine sleep disorders will be critical for clarifying this relationship. Such understanding could guide the development of targeted interventions ranging from management of BOAS to anti-inflammatory therapies that mitigate the cardiovascular risks of chronic sleep deprivation in dogs.

Hence In this study, a systematic meta-analysis of the existing scientific literature was undertaken to comprehensively evaluate the relationship between sleep deprivation and the modulation of inflammatory cytokines. The first phase of the investigation focused on identifying which cytokines most consistently exhibit altered expression in response to insufficient sleep across a variety of experimental and clinical settings. Through a structured search strategy, studies reporting quantitative measurements of cytokines in sleep-restricted or sleep-deprived subjects were collected, screened, and synthesised. Attention was given to cytokines frequently implicated in systemic inflammation, including members of the interleukin family and tumour necrosis factor pathways. Building upon these findings, the second phase of the study involved the construction of a domestic dog-specific cytokine interaction network to better understand how these sleep-sensitive cytokines may influence cardiac electrophysiology in dogs. Because the canine myocardium shares key physiological and molecular similarities with that of humans, this model provides valuable translational insight. Network analysis methods were employed to map interactions among the cytokines identified in the meta-analysis, integrating known signalling pathways, receptor relationships, and downstream molecular effects relevant to cardiac tissue. This systems-level approach enabled the identification of cytokine clusters and signalling hubs that may contribute to an increased susceptibility to arrhythmias. The integrated results of the meta-analysis and network modelling highlight how sleep deprivation induced shifts in inflammatory signalling could disrupt myocardial conduction through multiple interacting pathways. By delineating these cytokine associations within a domestic dog-specific context, the study provides a framework for understanding how chronic sleep loss may elevate arrhythmia risk and identifies potential molecular targets for future therapeutic or diagnostic efforts.

## Materials and methods

### Literature Review and Cytokine Selection

A targeted review of the scientific literature following PRISMA guidelines was conducted to identify cytokines reported to be associated with sleep deprivation. Two electronic databases (Google Scholar and PubMed) were systematically searched using the following key terms and Boolean combinations: “sleep loss” AND “cytokine storm”, and “chronic sleep deprivation” AND “inflammation”. The initial search retrieved a total of 25 articles. One duplicate record was removed, and the remaining 24 articles were screened for relevance based on predefined inclusion criteria, which required studies to report empirical measurements of inflammatory cytokines following experimentally induced or clinically observed sleep deprivation. Seven studies met all inclusion requirements and were selected for detailed analysis.^[26,31-36]^ From each study, data were extracted on cytokine expression patterns, measurement methodologies, subject characteristics, and the specific sleep-deprivation protocols used. Across these studies, seven cytokines were consistently identified as being modulated by sleep loss: interleukin-6 (IL6), interleukin-17A (IL17A), tumour necrosis factor-α (TNF), interleukin-1 (IL1), interleukin-21 (IL21), interferon-γ (IFNG), and C-reactive protein (CRP). These cytokines were selected as the basis for subsequent network and Gene Ontology enrichment analyses.

### Protein-Protein Interaction Network Construction

To investigate interaction patterns among the identified cytokines, a domestic dog-specific protein-protein interaction (PPI) network was generated using the STRING database (version 12.0). Each cytokine was queried individually and collectively in the STRING interface, with the organism parameter set to Canis lupus familiaris. The minimum interaction confidence score was set to 0.4 (medium confidence) unless otherwise indicated. Both experimentally validated interactions and computationally predicted associations were included to produce a comprehensive network model. The resulting interaction map was exported for visualization and further analysis.

### Gene Ontology (GO) Biological Process Enrichment Analysis

The set of seven cytokines was subjected to Gene Ontology (GO) Biological Process enrichment analysis using STRING’s built-in functional enrichment tools. The analysis identified biological processes significantly overrepresented among the cytokine group, using the STRING enrichment p-value (corrected for false-discovery rate) to determine statistical significance. Enriched processes related to inflammatory signalling, immune activation, and pathways previously implicated in cardiac electrophysiological modulation were prioritized for interpretation.

### Comparison of Major Cytokine Networks

The major interaction clusters and pathway associations identified in the STRING canine PPI network were compared qualitatively to highlight shared and unique interactions among the cytokines. Network topology features including degree centrality, interaction density, and functional clustering were examined to identify potential key regulators or hubs. These comparisons were used to evaluate how the collective cytokine network may contribute to inflammatory states associated with increased arrhythmia susceptibility in domestic dogs.

## RESULTS

The network analysis performed on the seven cytokines identified from the literature review (IL-6, IL-17A, TNF, IL-1, IL-21, IFN-γ, and CRP) revealed extensive molecular interactions that extended well beyond these initial markers (Figure 1). IL-6 and TNF appeared to be central within the network, each displaying multiple strong connections to the other cytokines. This centrality underscores their well-established roles as master regulators of systemic inflammation and suggests they may function as key integrators of the immune response to sleep loss. Their position as hubs aligns with the enrichment findings indicating activation of chemokine regulation, myeloid cell differentiation, and JAK-STAT signalling pathways. In contrast, CRP appeared more peripheral, with a single direct connection to IL-6. CRP is primarily a downstream acute-phase protein produced in response to IL-6 signalling, and its peripheral placement reflects this biology, rather than driving the inflammatory cascade, CRP serves as a biomarker of IL-6-mediated responses. Its connection to IL-6 suggests that elevations in CRP during sleep deprivation may be a consequence of upstream cytokine activation rather than a cause of broader network activity. When these cytokines were submitted to protein-protein interaction analysis, several additional molecules emerged as central hubs within the network. The most frequently involved and highly connected nodes included IL-10, IL-1β, JAK1, JAK2, IFN-γ, IL-6, STAT3, TNF, IL-2, and IL-4 (Figure 2). STAT3 was observed to be the key central hub for these networks. These molecules consistently appeared across multiple interaction maps and demonstrated strong connectivity patterns, suggesting that they play key regulatory roles in coordinating the inflammatory signalling cascades influenced by sleep deprivation. The prominence of JAK1, JAK2, and STAT3 within the network further indicated that signal transduction processes, particularly those involving JAK-STAT activation, may represent major convergence points for cytokine activity in the context of sleep loss (Figure 2).

**Figure 1:**
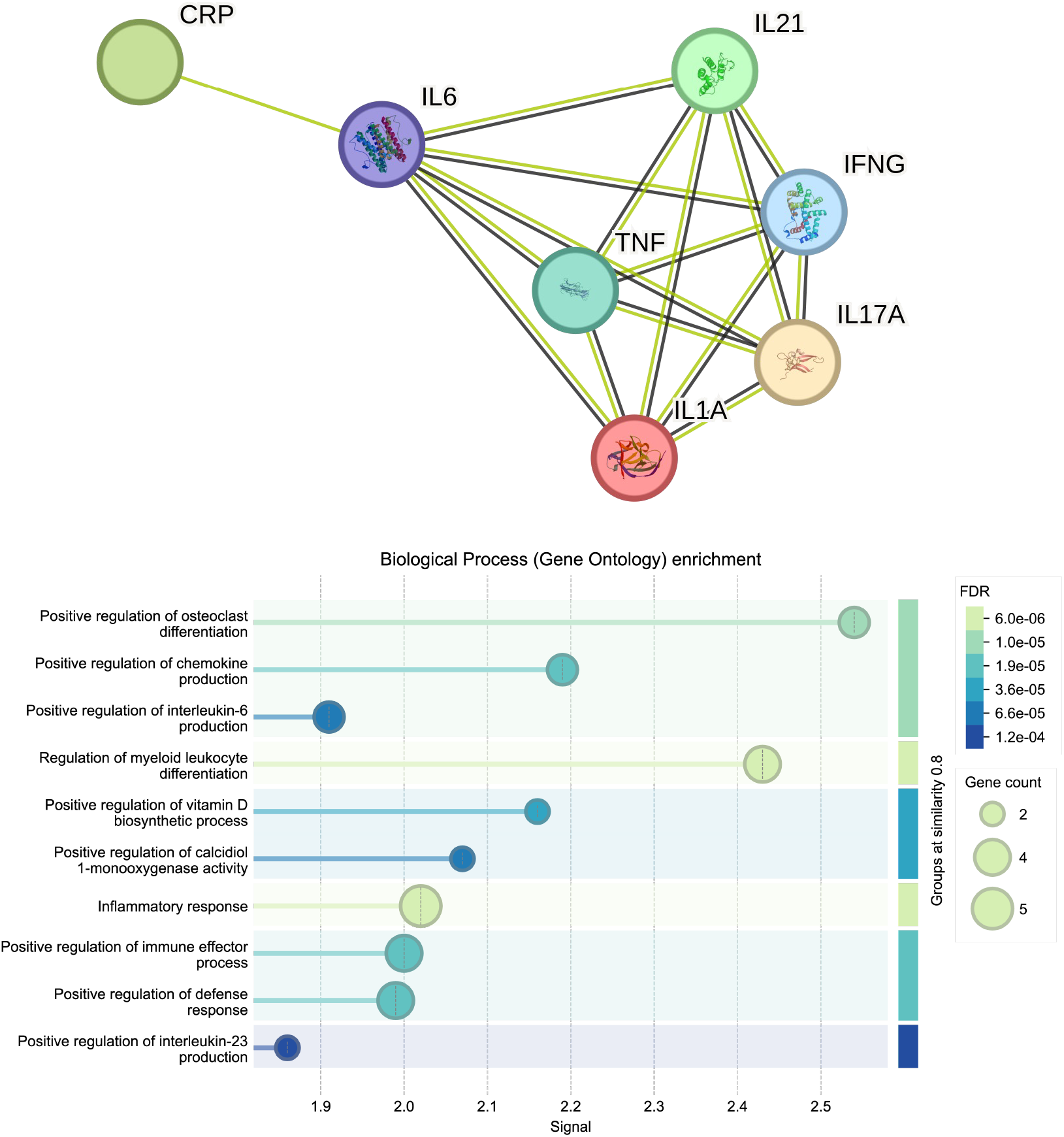
Interaction network and Gene Ontology (GO) Biological Process enrichment profile of the primary cytokines directly identified as associated with sleep deprivation, highlighting their core functional themes and molecular relationships.

**Figure 2:**
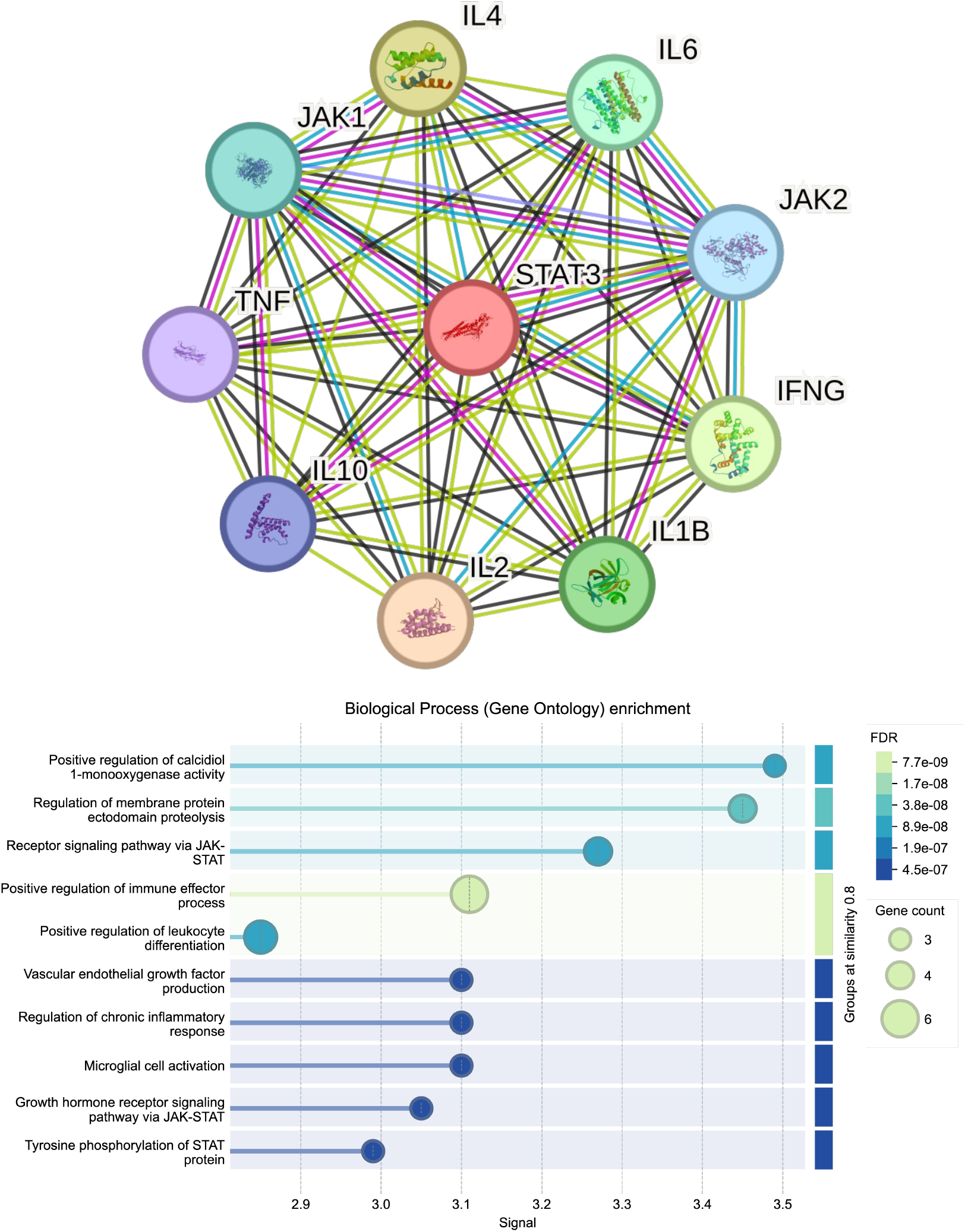
Interaction network and GO Biological Process enrichment analysis of the cytokines exhibiting the highest connectivity within the primary cytokine network, illustrating key hub molecules and the biological pathways they influence.

Primary enrichment analysis of this core network produced several statistically significant biological processes that were overrepresented among the interacting cytokines. The most notable processes included positive regulation of osteoclast differentiation, regulation of chemokine production, and positive regulation of myeloid leukocyte differentiation (Figure 1). The enrichment of osteoclast differentiation pathways suggests broader involvement of inflammatory mediators in bone-immune system interactions, which may be influenced by chronic inflammatory states. The regulation of chemokine production indicates heightened potential for immune-cell recruitment and trafficking, processes that are known to be sensitive to fluctuations in systemic inflammation. Additionally, the positive regulation of myeloid leukocyte differentiation reflects enhanced activation and maturation of innate immune cells, a pattern consistent with an organism experiencing physiological stress or perturbation, such as insufficient sleep.

Secondary enrichment analysis, which considered extended interactions surrounding the primary cytokine network, revealed additional processes of biological relevance. Among these, positive regulation of calcidiol 1-monooxygenase (CYP27B1) activity emerged as a significant finding. This enzyme plays an essential role in vitamin D metabolism, linking inflammatory signalling with endocrine regulatory pathways. The enrichment of membrane protein ectodomain proteolysis pointed to increased activity of proteolytic mechanisms involved in receptor remodelling, membrane instability and cytokine processing (Figure 2). Furthermore, signalling through the JAK-STAT pathway appeared strongly enriched within the extended network, reinforcing the initial observation that JAK1, JAK2, and STAT3 were among the most frequently involved signalling nodes (Figure 2). This consistency between network topology and enrichment analysis provides strong evidence that JAK-STAT signalling serves as a central mediator of the cytokine responses elevated during sleep deprivation.

When the primary and secondary networks were integrated into a comprehensive model, three major cytokine-cantered clusters became apparent (Figure 3). One cluster consisted predominantly of interleukin-related signalling, reflecting interactions among IL-6, IL-17A, IL-1, IL-21, and their associated downstream mediators. A second cluster cantered around CRP, representing pathways associated with the acute-phase response and systemic inflammatory modulation. The third prominent cluster was built around TNF and its extensive network of interactions, consistent with its well-established role as a master regulator of inflammatory activity. Collectively, the integrated network demonstrated a clear enrichment of cytokine-mediated signalling processes, indicating that the inflammatory effects of sleep deprivation operate through tightly interconnected pathways involving both classical pro-inflammatory cytokines and their downstream molecular partners. These findings highlight a coordinated biological response that may, in turn, contribute to downstream physiological disruptions, including increased susceptibility to arrhythmic events in the myocardium.

**Figure 3:**
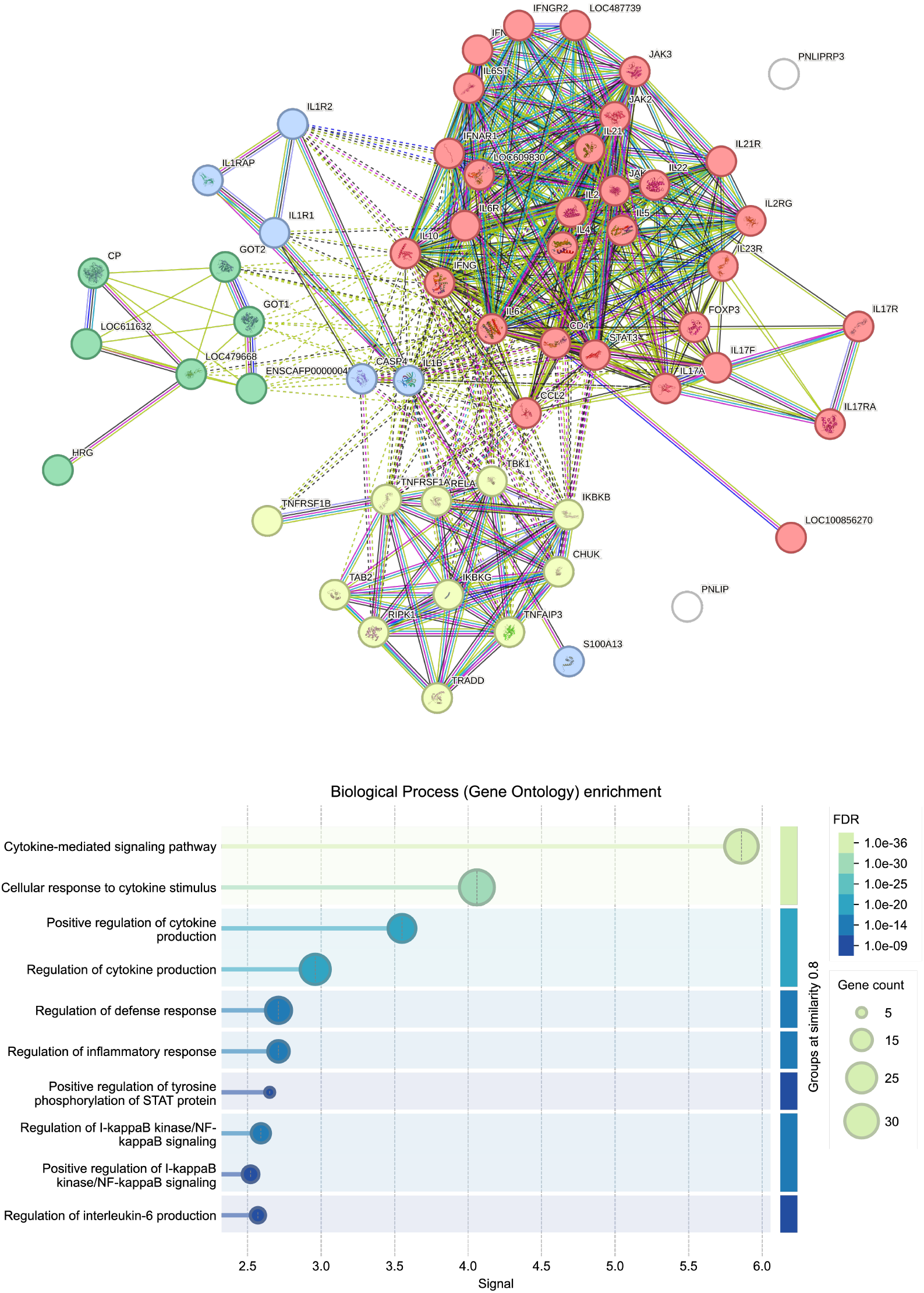
Comprehensive interaction network and GO Biological Process enrichment analysis of the full cytokine network associated with sleep deprivation, capturing both primary and secondary interactions and revealing broader functional pathway involvement.

## DISCUSSION

This study aimed to identify the cytokines associated with sleep deprivation and to explore, through network and enrichment analyses, how these inflammatory mediators may contribute to increased arrhythmia susceptibility in dogs. The findings reveal that sleep deprivation is linked to coordinated activation of multiple pro-inflammatory cytokines, with IL-6, TNF, IL-1A, IL-17A, IL-21, IFN-γ, and CRP forming a dense interaction network. The enrichment analyses identified several biological processes, including chemokine production, vitamin D metabolic regulation, positive osteoclast differentiation, membrane protein ectodomain proteolysis, and JAK-STAT signalling, that may mechanistically link sleep loss to myocardial electrical instability. Together, these pathways suggest a multifaceted inflammatory response with potential downstream effects on cardiac conduction and arrhythmogenesis.

One key finding of this study is the enrichment of pathways involved in the regulation of chemokine production. Chemokines play essential roles in recruiting immune cells to tissues and orchestrating inflammatory responses.^[37-39]^ Sleep deprivation has been shown in multiple studies to amplify chemokine expression, which may increase circulating leukocyte priming and their infiltration into myocardium.^[36,40]^ Immune-cell trafficking into myocardium can promote local inflammation, altering ion-channel expression, gap-junction integrity, and autonomic signalling.^[41,42]^ Such changes can reduce electrical conduction velocity and increase the dispersion of repolarisation, two hallmark contributors to arrhythmia vulnerability.^[43]^ The enrichment of vitamin D related metabolic processes, particularly positive regulation of calcidiol 1-monooxygenase (CYP27B1), suggests an additional mechanism linking sleep loss to inflammation and cardiac outcomes. CYP27B1 converts calcidiol to the active form of vitamin D, which has known immunomodulatory effects.^[44]^ Activated vitamin D typically suppresses pro-inflammatory cytokines and promotes regulatory immune phenotypes. An upregulation of CYP27B1-associated pathways under sleep-deprived conditions may reflect a compensatory response to inflammation.

However, sustained inflammatory activation may overwhelm this regulatory pathway, contributing to persistent cytokine elevation. Low or dysregulated vitamin D signalling has been associated with altered calcium homeostasis, impaired autonomic balance, and increased arrhythmic risk, providing another potential link between sleep deprivation and cardiac electrical instability.^[45,46]^

The identification of positive regulation of osteoclast differentiation in the enriched pathways further underscores the systemic impact of the sleep-deprivation induced cytokine milieu. Although osteoclast-related processes are typically associated with bone metabolism, they are tightly regulated by cytokines such as IL-6, IL-1, and TNF, which also exert potent cardiovascular effects.^[47,48]^ These cytokines can modify intracellular calcium handling, alter cardiomyocyte excitation-contraction coupling, and promote extracellular matrix remodelling within cardiac tissue.^[26,27,49]^ Such remodelling, even at a subtle level, can disrupt myocardial structural integrity and increase arrhythmia susceptibility by creating areas of non-uniform conduction or re-entry-prone circuits. Another enriched process, membrane protein ectodomain proteolysis, offers insight into how inflammatory signalling may influence cardiac electrophysiology. Ectodomain shedding is essential in regulating receptors, adhesion molecules, and ion-channel associated proteins at the cell surface. Inflammatory cytokines can activate proteases responsible for this shedding, potentially altering β-adrenergic receptor density, connexin turnover, and ion-channel surface expression on cardiomyocytes.^[50,51]^ These modifications can impair electrical coupling and destabilise conduction patterns, thereby enhancing vulnerability to arrhythmias.

The prominence of JAK-STAT signalling in the network further supports the view that sleep deprivation induces widespread pro-inflammatory signalling capable of influencing cardiac function. JAK-STAT activation by IL-6, IL-21, IFN-γ, and other cytokines leads to transcriptional changes that regulate cellular metabolism, immune-cell differentiation, and cytokine amplification loops.^[52-54]^ In the myocardium, chronic or excessive JAK-STAT activation can alter calcium-channel expression, disrupt mitochondrial function, and contribute to structural remodelling.^[52,54]^ These alterations are associated with increased risk of atrial and ventricular arrhythmias, especially under conditions of systemic inflammatory stress.^[55,56]^ The results of this study indicate that the cytokine network activated by sleep deprivation engages biological processes known to impact cardiac electrophysiology through inflammatory activation, ion-channel modulation, tissue remodelling, and autonomic dysregulation. Rather than acting through a single pathway, sleep deprivation appears to initiate a coordinated inflammatory response that converges on multiple mechanisms capable of altering myocardial conduction. These findings provide a mechanistic framework for understanding clinical observations linking insufficient sleep to elevated arrhythmia risk and highlight cytokine-driven pathways as potential targets for future investigation in both human and veterinary cardiac health.

Based on the mechanistic pathways identified in this analysis, it is likely that treating arrhythmias in sleep-deprived dogs requires more than the administration of anti-arrhythmic drugs alone. While anti-arrhythmics may provide temporary stabilisation of electrical conduction, the underlying drivers of myocardial irritability in this context appear to stem from a complex inflammatory environment characterized by elevated IL-6, TNF, IL-1 related signalling, chemokine upregulation, and JAK-STAT activation. Therefore, therapeutic strategies that target inflammation alongside arrhythmic control may offer greater clinical benefit.^[57,58]^ Potential approaches could include the use of anti-inflammatory agents, cytokine-modulating therapies, or interventions that help restore immune equilibrium, such as optimizing vitamin D status, reducing systemic inflammatory load, or supporting autonomic balance through stress-reduction and sleep-restorative measures. Modulating ectodomain proteolysis or osteoclast-linked inflammatory pathways may also hold translational promise, although such strategies remain experimental.^[59,60]^ Collectively, these findings suggest that addressing both the electrical instability and the inflammatory milieu is essential for fully managing arrhythmia risk in sleep-deprived dogs, and that a multimodal therapeutic strategy may be necessary to achieve sustained cardiac stabilization.

## Notes

### Competing Interest Statement

The authors have declared no competing interest.

## References

1. Devereux EA, Ejezie AV, Lynch AM, Gruen ME, LaJuett SJ, Robertson JB, Scharf VF. Factors Affecting Sleep Among Dogs and Cats in a Veterinary Intensive Care Unit. Journal of Veterinary Emergency and Critical Care. 2025:e13472.

2. Rudenko A, Rudenko P, Glamazdin I, Vatnikov Y, Kulikov E, Sachivkina N, Rudenko V, Sturov N, Babichev N, Romanova E. Assessment of Respiratory Rate in Dogs during the Sleep with Mitral Valve Endocardiosis, Complicated by Congestive Heart Failure Syndrome: the Degree of Adherence for this Test by Animal Owners and its Impact on Patient Survival. Systematic Reviews in Pharmacy. 2020;11.

3. Dumitrascu R, Heitmann J, Seeger W, Weissmann N, Schulz R. Obstructive sleep apnea, oxidative stress and cardiovascular disease: lessons from animal studies. Oxidative medicine and cellular longevity. 2013;2013:234631.

4. Mitze S, Barrs VR, Beatty JA, Hobi S, Bęczkowski PM. Brachycephalic obstructive airway syndrome: much more than a surgical problem. Veterinary Quarterly. 2022;42:213–223.

5. Ladlow J, Liu N-C, Kalmar L, Sargan D. Brachycephalic obstructive airway syndrome. The Veterinary Record. 2018;182:375.

6. Niinikoski I. Sleep-disordered breathing and inflammatory response in dogs. 2024.

7. Brown R, Pang G, Husband AJ, King MG, Bull DF. Sleep deprivation and the immune response to pathogenic and non-pathogenic antigens. Behavior and Immunity. 2019:127–134.

8. Besedovsky L, Lange T, Haack M. The sleep-immune crosstalk in health and disease. Physiological reviews. 2019.

9. Zhao J, Xu W, Yun F, Zhao H, Li W, Gong Y, Yuan Y, Yan S, Zhang S, Ding X. Chronic obstructive sleep apnea causes atrial remodeling in canines: mechanisms and implications. Basic research in cardiology. 2014;109:427.

10. Zhang Y-M, Wei R-M, Feng Y-Z, Zhang K-X, Ge Y-J, Kong X-Y, Li X-Y, Chen G-H. Sleep deprivation aggravates lipopolysaccharide-induced anxiety, depression and cognitive impairment: the role of pro-inflammatory cytokines and synaptic plasticity-associated proteins. Journal of neuroimmunology. 2024;386:578252.

11. Cui L, Xue R, Zhang X, Chen S, Wan Y, Wu W. Sleep deprivation inhibits proliferation of adult hippocampal neural progenitor cells by a mechanism involving IL-17 and p38 MAPK. Brain Research. 2019;1714:81–87.

12. Polidarová L, Houdek P, Sumová A. Chronic disruptions of circadian sleep regulation induce specific proinflammatory responses in the rat colon. Chronobiology international. 2017;34:1273–1287.

13. Al-Rashed F, Alsaeed H, Akhter N, Alabduljader H, Al-Mulla F, Ahmad R. Impact of sleep deprivation on monocyte subclasses and function. The Journal of Immunology. 2025;214:347–359.

14. Abedelmalek S, Chtourou H, Aloui A, Aouichaoui C, Souissi N, Tabka Z. Effect of time of day and partial sleep deprivation on plasma concentrations of IL-6 during a short-term maximal performance. European journal of applied physiology. 2013;113:241–248.

15. Chennaoui M, Sauvet F, Drogou C, Van Beers P, Langrume C, Guillard M, Gourby B, Bourrilhon C, Florence G, Gomez-Merino D. Effect of one night of sleep loss on changes in tumor necrosis factor alpha (TNF-α) levels in healthy men. Cytokine. 2011;56:318–324.

16. Helvig A, Wade S, Hunter-Eades L. Rest and the associated benefits in restorative sleep: a concept analysis. Journal of advanced nursing. 2016;72:62–72.

17. Jerath R, Harden K, Crawford M, Barnes VA, Jensen M. Role of cardiorespiratory synchronization and sleep physiology: effects on membrane potential in the restorative functions of sleep. Sleep medicine. 2014;15:279–288.

18. Milrad SF, Hall DL, Jutagir DR, Lattie EG, Ironson GH, Wohlgemuth W, Nunez MV, Garcia L, Czaja SJ, Perdomo DM. Poor sleep quality is associated with greater circulating pro-inflammatory cytokines and severity and frequency of chronic fatigue syndrome/myalgic encephalomyelitis (CFS/ME) symptoms in women. Journal of neuroimmunology. 2017;303:43–50.

19. Patel SR, Zhu X, Storfer-Isser A, Mehra R, Jenny NS, Tracy R, Redline S. Sleep duration and biomarkers of inflammation. Sleep. 2009;32:200–204.

20. Mesquita T, Lin YN, Ibrahim A. Chronic low-grade inflammation in heart failure with preserved ejection fraction. Aging Cell. 2021;20:e13453.

21. Kuusisto J, Kärjä V, Sipola P, Kholová I, Peuhkurinen K, Jääskeläinen P, Naukkarinen A, Ylä-Herttuala S, Punnonen K, Laakso M. Low-grade inflammation and the phenotypic expression of myocardial fibrosis in hypertrophic cardiomyopathy. Heart. 2012;98:1007–1013.

22. Irwin MR, Opp MR. Sleep health: reciprocal regulation of sleep and innate immunity. Neuropsychopharmacology. 2017;42:129–155.

23. Zielinski MR, Krueger JM. Sleep and innate immunity. Frontiers in bioscience (Scholar edition). 2011;3:632.

24. Wheeler ND, Ensminger DC, Rowe MM, Wriedt ZS, Ashley NT. Alpha-and beta-adrenergic receptors regulate inflammatory responses to acute and chronic sleep fragmentation in mice. PeerJ. 2021;9:e11616.

25. Lai C-T, Chen C-Y, Kuo TB, Chern C-M, Yang CC. Sympathetic hyperactivity, sleep fragmentation, and wake-related blood pressure surge during late-light sleep in spontaneously hypertensive rats. American journal of hypertension. 2016;29:590–597.

26. Kazanski V, Mitrokhin V, Mladenov M, Kamkin A. Cytokine effects on mechano-induced electrical activity in atrial myocardium. Immunological Investigations. 2017;46:22–37.

27. Kuzmin VS, Abramochkin DV, Mitrochin VM, Tian B, Makarenko EY, Kovalchuk LV, Khoreva MV, Nikonova A, Kalugin L, Lysenko NN. The role of proinflammatory cytokines in regulation of cardiac bioelectrical activity: link to mechanoelectrical feedback. Mechanical stretch and cytokines. 2011:107–153.

28. Oberholzer A, Oberholzer C, Moldawer LL. Cytokine signaling-regulation of the immune response in normal and critically ill states. Critical care medicine. 2000;28:N3-N12.

29. Belardelli F. Role of interferons and other cytokines in the regulation of the immune response. Apmis. 1995;103:161–179.

30. Krainer D, Dupré G. Brachycephalic obstructive airway syndrome. Veterinary Clinics: Small Animal Practice. 2022;52:749–780.

31. Kouvas N, Kontogiannis C, Georgiopoulos G, Spartalis M, Tsilimigras D, Spartalis E, Kapelouzou A, Kosmopoulos M, Chatzidou S. The complex crosstalk between inflammatory cytokines and ventricular arrhythmias. Cytokine. 2018;111:171–177.

32. Monden Y, Kubota T, Inoue T, Tsutsumi T, Kawano S, Ide T, Tsutsui H, Sunagawa K. Tumor necrosis factor-α is toxic via receptor 1 and protective via receptor 2 in a murine model of myocardial infarction. American Journal of Physiology-Heart and Circulatory Physiology. 2007;293:H743–H753.

33. Sang D, Lin K, Yang Y, Ran G, Li B, Chen C, Li Q, Ma Y, Lu L, Cui X-Y. Prolonged sleep deprivation induces a cytokine-storm-like syndrome in mammals. Cell. 2023;186:5500-5516. e5521.

34. Fontes JA, Rose NR, Čiháková D. The varying faces of IL-6: From cardiac protection to cardiac failure. Cytokine. 2015;74:62–68.

35. Tobaldini E, Costantino G, Solbiati M, Cogliati C, Kara T, Nobili L, Montano N. Sleep, sleep deprivation, autonomic nervous system and cardiovascular diseases. Neuroscience & Biobehavioral Reviews. 2017;74:321–329.

36. Irwin MR, Wang M, Campomayor CO, Collado-Hidalgo A, Cole S. Sleep deprivation and activation of morning levels of cellular and genomic markers of inflammation. Archives of internal medicine. 2006;166:1756–1762.

37. Harvanová G, Duranková S, Bernasovská J. The role of cytokines and chemokines in the inflammatory response. Alergologia Polska-Polish Journal of Allergology. 2023;10:210–219.

38. Palomino DCT, Marti LC. Chemokines and immunity. Einstein (são paulo). 2015;13:469–473.

39. Surmi BK, Hasty AH. The role of chemokines in recruitment of immune cells to the artery wall and adipose tissue. Vascular pharmacology. 2010;52:27–36.

40. Said EA, Al-Abri MA, Al-Saidi I, Al-Balushi MS, Al-Busaidi JZ, Al-Reesi I, Koh CY, Idris MA, Al-Jabri AA, Habbal O. Sleep deprivation alters neutrophil functions and levels of Th1-related chemokines and CD4+ T cells in the blood. Sleep and Breathing. 2019;23:1331–1339.

41. Mueller SN. Neural control of immune cell trafficking. Journal of Experimental Medicine. 2022;219:e20211604.

42. Nakai A, Leach S, Suzuki K. Control of immune cell trafficking through inter-organ communication. International Immunology. 2021;33:327–335.

43. Sanders E, Alcaide P. Red light-green light: T-cell trafficking in cardiac and vascular inflammation. American Journal of Physiology-Cell Physiology. 2022.

44. Anderson PH, O’Loughlin PD, May BK, Morris HA. Modulation of CYP27B1 and CYP24 mRNA expression in bone is independent of circulating 1, 25 (OH) 2D3 levels. Bone. 2005;36:654–662.

45. Barsan M, Brata AM, Ismaiel A, Dumitrascu DI, Badulescu A-V, Duse TA, Dascalescu S, Popa SL, Grad S, Muresan L. The pathogenesis of cardiac arrhythmias in vitamin D deficiency. Biomedicines. 2022;10:1239.

46. Demir M, Uyan U, Melek M. The effects of vitamin D deficiency on atrial fibrillation. Clinical and applied thrombosis/hemostasis. 2014;20:98–103.

47. Salari P, Abdollahi M. A comprehensive review of the shared roles of inflammatory cytokines in osteoporosis and cardiovascular diseases as two common old people problem; actions toward development of new drugs. International Journal of Pharmacology. 2011;7:552–567.

48. Turner NA, Mughal RS, Warburton P, O’Regan DJ, Ball SG, Porter KE. Mechanism of TNFα-induced IL-1α, IL-1β and IL-6 expression in human cardiac fibroblasts: effects of statins and thiazolidinediones. Cardiovascular research. 2007;76:81–90.

49. Sugishita K, Kinugawa K-i, Shimizu T, Harada K, Matsui H, Takahashi T, Serizawa T, Kohmoto O. Cellular basis for the acute inhibitory effects of IL-6 and TNF-α on excitation-contraction coupling. Journal of molecular and cellular cardiology. 1999;31:1457–1467.

50. Lundby A, Andersen MN, Steffensen AB, Horn H, Kelstrup CD, Francavilla C, Jensen LJ, Schmitt N, Thomsen MB, Olsen JV. In vivo phosphoproteomics analysis reveals the cardiac targets of β-adrenergic receptor signaling. Science signaling. 2013;6:rs11–rs11.

51. Rodrigues SF, Tran ED, Fortes ZB, Schmid-Schönbein GW. Matrix metalloproteinases cleave the β2-adrenergic receptor in spontaneously hypertensive rats. American Journal of Physiology-Heart and Circulatory Physiology. 2010;299:H25–H35.

52. Wu J-W, Wang B-X, Shen L-P, Chen Y-L, Du Z-Y, Du S-Q, Lu X-J, Zhao X-D. Investigating the Potential Therapeutic Targeting of the JAK-STAT Pathway in Cerebrovascular Diseases: Opportunities and Challenges. Molecular Neurobiology. 2025:1–27.

53. Lv Y, Qi J, Babon JJ, Cao L, Fan G, Lang J, Zhang J, Mi P, Kobe B, Wang F. The JAK-STAT pathway: from structural biology to cytokine engineering. Signal transduction and targeted therapy. 2024;9:221.

54. Parra-Izquierdo I, Sánchez-Bayuela T, Castaños-Mollor I, López J, Gómez C, San Román JA, Sanchez Crespo M, García-Rodríguez C. Clinically used JAK inhibitor blunts dsRNA-induced inflammation and calcification in aortic valve interstitial cells. The FEBS Journal. 2021;288:6528–6542.

55. Jiang H, Yang J, Li T, Wang X, Fan Z, Ye Q, Du Y. JAK/STAT3 signaling in cardiac fibrosis: A promising therapeutic target. Frontiers in Pharmacology. 2024;15:1336102.

56. Patel NJ, Nassal DM, Gratz D, Hund TJ. Emerging therapeutic targets for cardiac arrhythmias: Role of STAT3 in regulating cardiac fibroblast function. Expert opinion on therapeutic targets. 2021;25:63–73.

57. Dobrev D, Heijman J, Hiram R, Li N, Nattel S. Inflammatory signalling in atrial cardiomyocytes: a novel unifying principle in atrial fibrillation pathophysiology. Nature Reviews Cardiology. 2023;20:145–167.

58. Armbruster AL, Campbell KB, Kahanda MG, Cuculich PS. The role of inflammation in the pathogenesis and treatment of arrhythmias. Pharmacotherapy: The Journal of Human Pharmacology and Drug Therapy. 2022;42:250–262.

59. Wang Y, Zhong Z, Ma M, Zhao Y, Zhang C, Qian Z, Wang B. The role played by ailanthone in inhibiting bone metastasis of breast cancer by regulating tumor-bone microenvironment through the RANKL-dependent pathway. Frontiers in Pharmacology. 2023;13:1081978.

60. Mead TJ, Bhutada S, Martin DR, Apte SS. Proteolysis: a key post-translational modification regulating proteoglycans. American Journal of Physiology-Cell Physiology. 2022;323:C651–C665.

